# Single-base mapping of m6A in lncRNAs reveals a distinct landscape linked to transposable elements and RNA processing

**DOI:** 10.64898/2026.10.30.762970

**Authors:** Euijin Kwon, Zixiu Li, Chan Zhou

## Abstract

N6-methyladenosine (m6A) is the most abundant internal RNA modification with established roles in mRNA metabolism. Although m6A has been characterized in several well-known long noncoding RNAs (lncRNAs), its transcriptome-wide landscape and roles remain poorly understood. Here, we characterized the m6A landscape of lncRNAs at single-base resolution, identified m6A deposition-associated features, and investigated potential roles of m6A in lncRNA processing. Unlike in mRNAs, m6A sites are broadly distributed along lncRNAs. We further found that these differences in m6A patterns between lncRNAs and mRNAs may reflect contributions from transposable element-derived sequence to motif availability and local RNA structure, together with splicing-related constraints on m6A deposition near exon junctions. Additionally, m6A may be relevant to the regulation of lncRNA polyadenylation. Together, these findings define the single-base-resolution m6A landscape in lncRNAs, and connect lncRNA methylation to transposable elements, RNA structure, splicing, and polyadenylation. This study broadens our understanding of m6A in the lncRNA epitranscriptome.

## INTRODUCTION

RNA modifications constitute an important layer of post-transcriptional regulation to gene expression.^1–3^ To date, more than 170 distinct RNA modifications have been identified in all classes of RNA molecule in living organism. Among them, N6-methyladenosine (m6A) is known to be the most prevalent internal modification in eukaryotic RNA.^4,5^ m6A, a reversible modification, is installed by m6A writer complexes, removed by m6A demethylases, and recognized by m6A reader proteins. Through interacting with m6A reader proteins, m6A regulates multiple aspect of mRNA metabolisms, including splicing, stability, and translation.^6–12^ These regulatory roles have broad implications for biological processes and human disease. For example, m6A has been shown to modulate cell differentiation, embryonic development and spermatogenesis^13–16^, while its dysregulation has been linked to cancer and neurodevelopmental disease.^17–20^

In addition to mRNAs, m6A modifications have also been found in several long noncoding RNAs (lncRNAs).^21–24^ LncRNAs are conventionally defined as RNA transcripts longer than 200 nucleotides that lack protein-coding potential.^25,26^ lncRNAs participate in diverse cellular processes by regulating gene expression at the transcriptional, and post-transcriptional level, including by modulating chromatin architecture, transcription, RNA splicing, stability, localization and translation.^25,27^ Many lncRNAs undergo splicing and 3′-end cleavage and polyadenylation, which shape their transcript isoforms and expression.^25^ LncRNAs also have distinct sequence and structural features; for example, many contain transposable element-derived sequences that contribute to their diversity.^28^ These properties raise the possibility that m6A deposition in lncRNAs may have a landscape and regulatory context distinct from those in mRNAs.

Although m6A has been extensively characterized in mRNA, the transcriptome-wide m6A landscape and the factors associated with its deposition in lncRNA remain poorly understood. Many commonly used antibody-enrichment methods, including MeRIP-seq identify m6A-enriched regions rather than the precise modified nucleotide.^29^ This regional resolution limits the ability to relate individual m6A sites to local sequence or structural features or specific RNA-processing events. Mapping m6A at single-base resolution enables more precise analysis of these relationships, including splicing and 3′-end formation.

Here, we performed an integrative multi-omics analysis to systematically characterize the m6A landscape in lncRNAs at single-base resolution and investigate its potential relationship to lncRNA processing. We identified a distinct m6A landscape in lncRNAs compared with mRNAs and found that m6A deposition was associated with transposable element-derived sequences and structural features. Across lncRNAs, m6A deposition was inversely associated with transcript exon number. Moreover, depletion of the exon junction complex (EJC) increased the number of m6A-containing internal exons, suggesting that the EJC may constrain m6A deposition during lncRNA splicing. Finally, m6A was enriched near non-canonical lncRNA cleavage sites and associated with alternative polyadenylation usage. These findings define a site-resolved m6A landscape in lncRNAs and connect m6A with transposable elements and key aspects of lncRNA processing.

## RESULTS

### lncRNAs exhibit higher m6A level and m6A site frequency than mRNAs

To examine overall m6A levels between lncRNAs and mRNAs, we analyzed m6A-LAIC-seq (m6A-level and isoform-characterization sequencing) data, which separately profile m6A-positive (eluate) and m6A-negative (supernatant) RNA fractions separately (**Figure 1A)**.^30^ Using m6A-LAIC-seq data from GM12878 and H1 ESCs, we quantified m6A levels for expressed lncRNA and mRNA gene (**Table S2)**. The distribution of m6A levels showed that lncRNA genes tend to have higher m6A level compared to mRNAs genes in both cell lines (**Figure 1B** and **Figure S1A)**.

**Figure 1.**
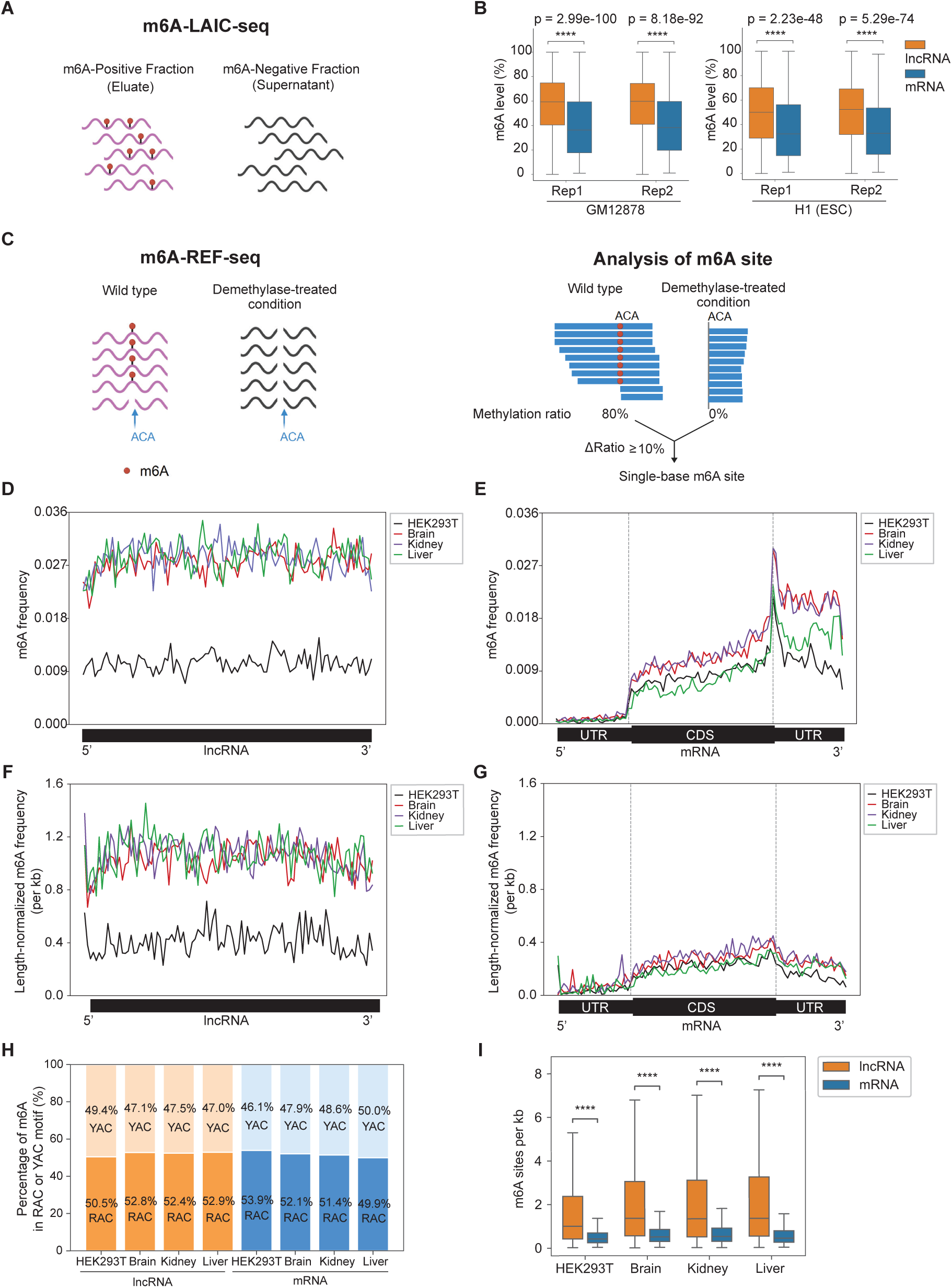
**Single-base-resolution mapping reveals distinct m6A landscapes in lncRNAs and mRNAs** (A) Schematic of m6A-LAIC-seq, which separates m6A-positive RNA (eluate) from m6A-negative RNA (supernatant). (B) m6A levels stratified by types of RNA. Box plots comparing m6A levels of different classes of RNA (lncRNA vs. mRNA genes) for GM12878 (left) and H1 ESC cells (right), p-values are calculated by two-sided Wilcoxon test. (C) Left, schematic diagram of m6A-REF-seq data. RNA from untreated wild-type and FTO-treated samples is exposed to MazF, which cleaves unmethylated ACA motifs; m6A at an ACA motif protects it from cleavage. Untreated wild-type samples therefore contain both cleaved and uncleaved molecules, whereas FTO-mediated demethylation makes previously protected ACA motifs susceptible to cleavage. Right, schematic of single-base m6A-site calling. For each ACA motif, the ratio of reads containing the motif internally to reads terminating at the motif is calculated in each condition. Sites with a difference in this ratio of >10% between untreated and FTO-treated samples are called m6A sites. (D) Transcriptome-wide distribution of m6A frequency across lncRNA transcripts in HEK293T cells and multiple human tissues. Each lncRNA transcript was divided into 100 equal-length bins. m6A frequency was calculated as the number of m6A sites per bin normalized to the number of expressed lncRNA transcripts. (E) Transcriptome-wide distribution of m6A frequency across mRNA transcripts in HEK293T cells and multiple human tissues. The 5’ UTR, CDS, 3’ UTR were divided into 25, 50, 25 equal-length bins, respectively. m6A frequency was calculated as the number of m6A sites per bin to the number of expressed transcripts containing the corresponding region. (F) Transcriptome-wide length-normalized m6A frequency per kilobase across lncRNA transcripts. Transcripts were binned as in (D), and m6A site counts in each bin were normalized by the corresponding sequence length and the number of expressed lncRNA transcripts and multiplied by 1,000. (G) Transcriptome-wide length-normalized m6A frequency per kilobase across mRNA transcripts. Transcript regions were binned as in (E), and m6A site counts in each bin were normalized by the corresponding regional sequence length and the number of expressed transcripts containing that region and multiplied by 1,000. (H) Proportions of RAC and YAC m6A sub-motifs in lncRNAs and mRNAs. R denotes A or G; Y denotes C or U. (I) Number of m6A sites per kilobase of transcript length in lncRNAs and mRNAs, calculated separately for each transcript. **** denotes p<= 1.00e-04 which were calculated with Mann-Whitney-Wilcoxon two-sided test.

To further characterize the m6A landscape in lncRNA transcriptome at single-base resolution, we analyzed m6A-REF-seq (m6A-sensitive RNA-Endoribonuclease–Facilitated sequencing) data (**Figure 1C, Left**).^31^ By comparing the wild type with m6A modification and demethylated condition, we identified single-base m6A sites in lncRNAs and mRNAs **(Figure 1C, Right; Table S3).** We first examined transcriptome-wide m6A frequency across transcripts expressed in HEK293T cells and multiple human tissues. lncRNAs showed a relatively even distribution of m6A frequency along their length from the 5’ to 3’ end, whereas mRNAs exhibited prominent enrichment near the CDS–3′ UTR boundary and within the 3′ UTR, consistent with previous studies **(Figure 1D and 1E)**.^29,31^

Because differences in transcript and regional sequence length influence these profiles, we next examined length-normalized m6A distributions. After accounting for sequence length, lncRNAs retained a relatively uniform m6A profile **(Figure 1F)**. In contrast, mRNAs showed a gradual increase in m6A frequency toward the end of CDS, with only modest enrichment near the CDS-3’UTR boundary followed by a decline across the 3’UTR **(Figure 1G)**. The reduced enrichment after length normalization indicates that differences in regional sequence length partially contribute to the prominent peak observed in the non-length-normalized profile.

To determine whether the transcriptome-wide profiles reflected broadly distributed m6A sites or enrichment within a subset of transcripts, we next examined m6A patterns at the individual-transcript level. Heatmap analysis revealed dispersed m6A enrichment across many lncRNA transcripts, collectively producing the relatively uniform transcriptome-wide profile. In contrast, fewer than 20% of mRNA transcripts contained m6A sites near the CDS-3’UTR boundary, indicating that alignment of site enrichment within a subset of transcripts can contribute to the prominent aggregate peak observed in the transcriptome-wide mRNA profile **(Figure S1B)**.

We next asked whether the distinct m6A distribution patterns between lncRNAs and mRNAs were influenced by the different positional framework used for the two RNA classes. We therefore stratified transcripts by exon number and applied the same exon-based positional framework to both lncRNA and mRNAs. m6A density was calculated by normalizing m6A site counts by both bin length and the number of transcripts contributing to each region. Single-exon, two-exon, and ≥3-exon lncRNAs represented 25.6% and 43.0%, 31.4% of the analyzed lncRNA transcripts, respectively, and showed relatively uniform m6A distributions with only modest regional variation **(Figure S1C).** By contrast, ≥3-exon mRNAs, which accounted 89.8% of the analyzed mRNA transcripts, displayed a more pronounced positional pattern, with m6A frequency increasing toward the end of the internal exon and subsequently declining across the last exon **(Figure S1D)**. Broadly similar patterns were observed in human brain, kidney, and liver samples **(Figure S1E and S1F)**. Fold change analysis of mean regional m6A density further showed that increases from the first region/exon toward internal and terminal regions/exon were modest in lncRNAs but substantially more pronounced in ≥3-exon mRNAs (**Figure S1G and S1H)**. Overall, lncRNAs exhibited even m6A distribution, whereas mRNAs showed more pronounced regional enrichment, even when both RNA classes were analyzed using the same exon-based positional framework.

Canonical m6A motifs typically contain a purine immediately upstream of the modified adenosine, forming an RAC motif. However, approximately half of the m6A sites detected in our dataset were located within RAC motif, whereas the remainder occurred in YAC motif (Y=C/U). To examine whether motif preference differed between lncRNAs and mRNAs, we compared the proportions of RAC-and YAC-associated m6A sites in the two transcript classes. RAC and YAC proportions were comparable between lncRNAs and mRNAs, with RAC motifs slightly more prevalent in both groups **(Figure 1H)**.

Finally, to compare the overall density of m6A sites while accounting for transcript length, we calculated the number of detected m6A sites per kilobase of individual transcript sequence. lncRNAs exhibited higher m6A site density than mRNAs in HEK293T cells and across the examined human tissues **(Figure 1I)**. Together, these analyses indicate that lncRNAs exhibit higher m6A levels and greater length-normalized m6A site frequency than mRNAs, while showing less pronounced positional enrichment across transcript regions.

### Transposable elements-derived sequences and local RNA structure are associated with distinct m6A deposition context

Previous study demonstrated that lncRNA contains a substantial fraction of exonized TE sequence. Majority of human lncRNA transcripts contain an exon of at least partial TE sequence, and about one third of total length of intergenic lncRNAs are covered by TE sequence. mRNA exons are also known have TE sequence, but showing the significant regional preference in 3’UTR compared to 5’UTR and CDS regions.^28^ Considering the enrichment of TE in lncRNA and 3’UTR mRNA are consistent with the m6A enrichment in the same region, we hypothesized that TE may be associated with the m6A deposition.

We first characterized the transcriptome-wide distribution of TEs in lncRNA and mRNA. TE-derived sequences were broadly distributed across lncRNA transcripts, with a modest increase toward the 3’ end, whereas mRNAs showed pronounced TE enrichment within the 3’UTR **(Figure 2A)**. These overall patterns were retained after normalization for regional sequence length **(Figure 2B)**.

**Figure 2.**
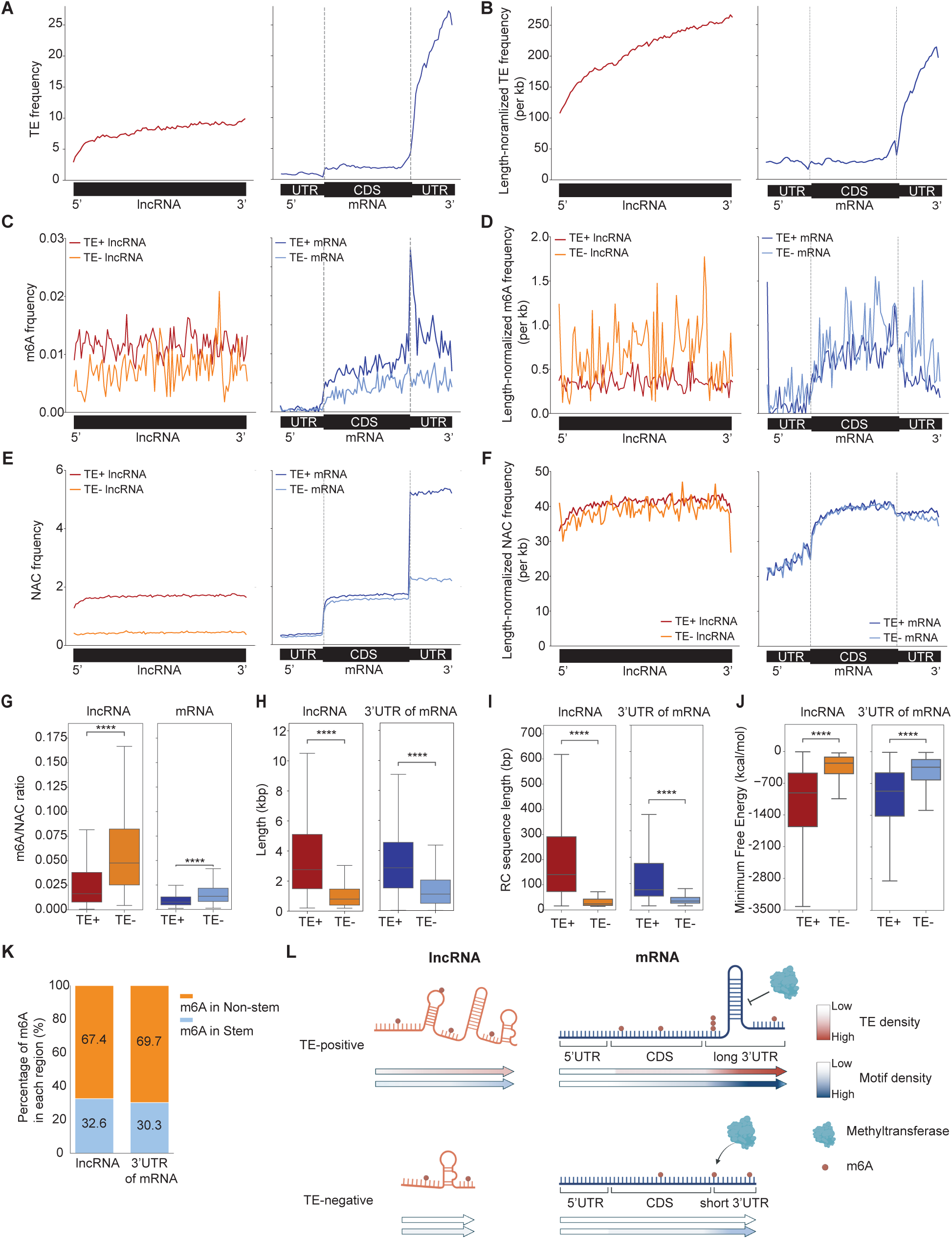
Transposable element-derived sequences and local RNA structure may contribute to distinct m6A patterns between lncRNAs and mRNAs. (A) Transcriptome-wide distribution of transposable elements (TEs) across lncRNAs and mRNAs expressed in HEK293T cells. Each lncRNA transcript was divided into 100 equal-length bins. For mRNAs, the 5′ UTR, CDS, and 3′ UTR were divided into 25, 50, and 25 equal-length bins, respectively. The number of TE-overlapping nucleotides in each bin was normalized to the total number of expressed transcripts containing the corresponding region. (B) Transcriptome-wide distribution of length-normalized TE density per kilobase across lncRNAs and mRNAs expressed in HEK293T cells. lncRNA and mRNA transcripts were partitioned into bins as described in (A). For each bin, the number of TE-overlapping nucleotides was normalized by the corresponding sequence length and by the number of expressed transcripts containing that region. (C) Distribution of m6A frequency across TE-positive (TE+) and TE-negative (TE–) lncRNAs and mRNAs. (D) Transcriptome-wide distribution of length-normalized m6A frequency per kilobase across TE-positive (TE+) and TE-negative (TE–) lncRNAs and mRNAs. (E) Transcriptome-wide distribution of NAC motif frequency across TE-positive and TE-negative lncRNAs and mRNAs. N denotes any nucleotide. (F) Transcriptome-wide distribution of length-normalized NAC motif frequency per kilobase across TE-positive and TE-negative lncRNAs and mRNAs. N denotes any nucleotide (G) m6A/NAC ratios in TE-positive and TE-negative lncRNAs and mRNAs. (H) Length distributions of TE-positive and TE-negative lncRNAs and 3′ UTR of mRNAs. (I) Lengths of reverse-complementary (RC) sequences identified by BLASTN in TE-positive and TE-negative lncRNAs and 3′ UTR of mRNAs. (J) Minimum free energy (MFE) of TE-positive and TE-negative lncRNAs and 3′ UTR of mRNAs. A lower MFE value indicates a more stable secondary structure. (K) Length-normalized proportion of m6A sites in stem and non-stem regions of lncRNAs and 3′ UTR of mRNAs. (L) Proposed model illustrating how TE-derived sequences may contribute to m6A deposition. (C-F): Darker and lighter lines represent TE-positive and TE-negative transcripts, respectively. Statistical test: From (G) to (J), ****p ≤ 1 × 10⁻⁴ by two-sided Mann–Whitney U tests.

We next examined whether the presence of TEs was associated with m6A distribution by classifying transcripts as TE-positive (TE+) or TE-negative (TE−) **(Table S4).** For non-length normalized m6A frequency, both TE+ and TE-lncRNAs showed broadly distributed m6A patterns without pronounced positional enrichment, although TE+ lncRNAs shows higher than TE-lncRNAs **(Figure 2C and S2A).** In mRNA, TE-transcripts showed reduced m6A frequency and lacked the prominent enrichment near the CDS-3’UTR boundary observed in TE+ transcripts **(Figure 2C and S2B)**. However, after normalization for regional sequence length, TE-lncRNAs and mRNAs tended to show higher m6A frequency than their TE+ counterparts, although substantial variation was observed **(Figure 2D)**.

We next investigated whether differences in NAC motif availability were associated with these TE-dependent patterns. For non-length-normalized NAC motif frequency, TE+ lncRNAs showed higher values than TE-lncRNAs across transcript regions, whereas in mRNAs, the difference between TE+ and TE-transcripts was most pronounced in the 3’ UTR, where TE-derived sequences were preferentially enriched. **(Figure 2E and S2C, 2SD)**. After normalization for regional sequence length, the differences in NAC motif frequency between TE+ and TE− groups were substantially reduced **(Figure 2F)**. To more directly assess the contribution of TE-derived sequences themselves, we compared NAC motif density between TE-overlapping and non-TE regions. NAC motif density was significantly higher within TE-overlapping regions, indicating that TE-derived sequences provide NAC-rich contexts that may increase the local availability of potential m6A motifs **(Figure S2E)**.

Despite the greater availability of NAC motifs in TE+ transcripts, the fraction of available motifs associated with detected m6A sites showed the opposite pattern. The m6A/NAC ratio was significantly lower in TE+ than in TE− lncRNAs and mRNAs **(Figure 2G and Figure S2F)**. This discrepancy was also evident in the positional profiles of TE+ mRNAs: although NAC motif frequency remained elevated within the 3′ UTR, m6A frequency declined more sharply across this region. Together, these observations indicate that motif availability alone is insufficient to explain m6A deposition in TE-containing transcripts and suggest that additional features may influence whether an available motif is methylated.

We therefore asked whether sequence and structural features associated with TE-containing transcripts could provide an additional constraint on m6A deposition. TE+ lncRNAs and TE+ mRNA 3′ UTRs were significantly longer than their TE− counterparts **(Figure 2H and Figure S2G)**, potentially providing greater sequence opportunity for intramolecular interactions. Consistent with this possibility, TE+ lncRNAs and mRNA 3′ UTRs contained longer reverse-complementary (RC) sequence matches than their TE− counterparts **(Figure 2I)**. RNA secondary-structure prediction further revealed differences in predicted minimum free energy between TE+ and TE− sequences **(Figure 2J and Figure S2H)**. Within predicted TE+ RNA structures, length-normalized m6A sites were preferentially located in non-stem regions relative to stem regions **(Figure 2K and Figure S2I)**, supporting an association between local structural accessibility and m6A deposition.

Together, these analyses highlight the importance of accounting for regional sequence length when interpreting TE-associated differences in m6A frequency. They further support a model in which TE-derived sequence increase the availability of potential m6A motifs through local enrichment of NAC motif. At the same time, the repetitive sequence context of TEs may promote RNA secondary structure and thereby reduce the accessibly of a subset of motifs for methylation **(Figure 2L)**. Thus, TE-derived sequences may influence the lncRNA and mRNA m6A landscapes through a balance between increased motif availability and local structural constraints.

### m6A deposition in lncRNAs decreases with exon number and may be restrained by exon junction complex

Previous studies found the negative relationship between mRNA splicing and m6A deposition in mRNAs and suggested splicing event may inhibit m6A deposition.^32^ Because lncRNA transcript tend to have a smaller number of exons compared to mRNA genes^33^, we hypothesized that lncRNAs may have higher m6A deposition rate compared to mRNAs. To test this, we examined the m6A/NAC ratio in lncRNAs and mRNAs and found that, as expected, lncRNAs had a higher m6A/NAC ratio than mRNAs; this difference was reproduced in brain, kidney, and liver. **(Figure 3A and S3A).** Within lncRNAs, the ratio decreased with increasing exon number in HEK293 cells (**Figure 3B**; slope = −4.65 × 10^−3^; p = 3.00 × 10^−10^). The association was consistent across human tissues, with slopes of −4.84 × 10^−3^ in brain, −6.26 × 10^−3^ in kidney, and −5.42 × 10^−3^ in liver (**Figure S3B**). Thus, lncRNAs with fewer annotated exons carried more detectable m6A sites relative to NAC content, although exon number here reflects transcript architecture, rather than a direct measurement of splicing efficiency.

**Figure 3.**
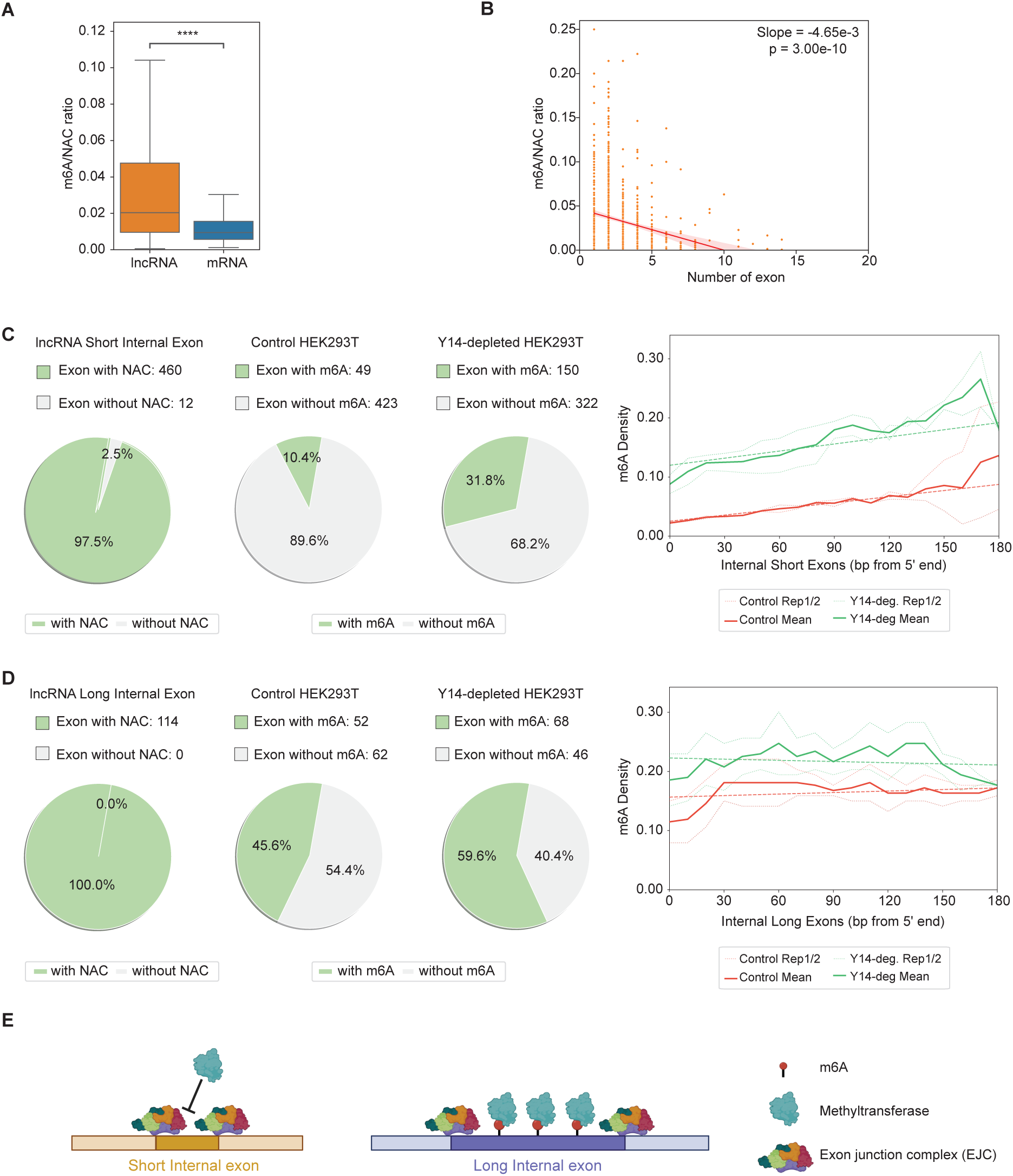
m6A deposition in lncRNA is negatively correlated with exon number and is sensitive to EJC near exon junctions. (A) m6A/NAC ratio in lncRNA and mRNAs. **** denotes p<= 1.00e-04 which were calculated with Mann-Whitney-Wilcoxon two-sided test. (B) The scatter plot showing the relationship m6A/NAC ratio and exon numbers for individual lncRNA transcripts. Slope and p value were calculated using the linear regression model. (C-D) Analysis of short (≤200 nt; C) and long (>200 nt; D) lncRNA internal exons. Pie charts show the numbers and proportions of exons with or without NAC motifs (left), with or without m6A in control HEK293T cells (middle), and with or without m6A after Y14 depletion (right). Line plots show m6A density by position along the internal exon, measured from the 5′ end, in control cells (red) and Y14-depleted cells (green). Replicates and group means are shown. (E) Proposed model in which EJC occupancy near exon-intron boundaries limits m6A deposition in lncRNA internal exons.

The exon junction complex (EJC) can restrict methylation near splice junctions in mRNAs, through blocking m6A methyltransferase to access the exon-intron boundary during mRNA splicing.^34–36^ Hence, to determine whether EJC can also restrict m6A methylation near splice junctions in lncRNAs, we reanalyzed published HEK293T m6A immunoprecipitation data following depletion of the EJC component Y14^35^ for both short (≤ 200 nt) and long (>200 nt) internal exons in lncRNAs **(Table S4)**. Because these data are peak-based rather than single-base, we scored each exon for the presence or absence of m6A. Nearly all short internal exons (≤200 nt) contained at least one NAC motif (460 of 472; 97.5%), yet only 10.4% (49 of 472) carried m6A in control cells. Following Y14 depletion, the proportion of m6A peak-containing internal exon increased significantly from 10.4% to 31.8% (150 of 472; p = 3.21 × 10⁻²², two-sided exact McNemar’s test), and m6A density increased along the entire length of short exons rather than at a single position (**Figure 3C**). Both replicate profiles showed higher m6A-enrichment ratio across 5’ end of short internal exons after Y14 depletion. Long internal exons (>200 nt), all of which contained NAC motifs, were already frequently methylated in control cells (45.6%; 52 of 114) and showed a smaller relative increase upon Y14 depletion (59.6%; 68 of 114; p = 0.00372, two-sided exact McNemar’s test), together with a modest increase in positional density **(Figure 3D; Table S5)**. The greater increase in m6A methylation in short than in long internal exons following Y14 depletion is consistent with an EJC-associated protective effect that extends across short exons but is more restricted to junction-proximal regions in longer exons **(Figure 3E).** However, most short internal exons (68.2%) remained unmethylated after Y14 depletion, suggesting that EJC-associated protection alone does not fully explain the low methylation of lncRNA internal exons, as previously reported for mRNAs.^32^ Together, these findings support a role for the EJC in limiting m6A deposition in lncRNA internal exons and are consistent with the possibility that the lower exon number of lncRNAs contributes to their higher methylation efficiency.

### m6A is associated with 3′ end cleavage-site context and alternative polyadenylation in a subset of lncRNAs

Polyadenylation process is crucial post-transcriptional processes that involve an endonucleolytic cleavage followed by the addition of a chain of adenine nucleotides. This process is regulated by cleavage and polyadenylation machinery that target a cis-regulatory RNA motif, polyA signal (PAS) located in 11-24nt upstream of cleavage site and GU/U-rich element **(Figure 4A)**.^37^ Multiple PAS allows the generation of different isoforms through alternative polyadenylation (APA). In mRNAs, m6A has been linked to 3′-end processing: methylated isoforms preferentially use proximal poly(A) sites^30^, m6A is enriched at non-canonical cleavage sites in human tissues,^38^ and the methyltransferase component and m6A reader have been implicated in APA.^39,40^ However, whether m6A is associated with lncRNA 3’-end formation has not been examined, while 3′-end formation of lncRNAs differs from that of mRNAs: lncRNAs may terminate transcription independently of the polyadenylation signal (PAS) and are less efficiently polyadenylated.^41^ We therefore assembled a catalog of cleavage sites that combines PolyA_DB annotations^42^ with sites inferred from RNA-seq coverage (**Table S6; STAR Methods**) and examined the distribution of m6A around these sites.

**Figure 4.**
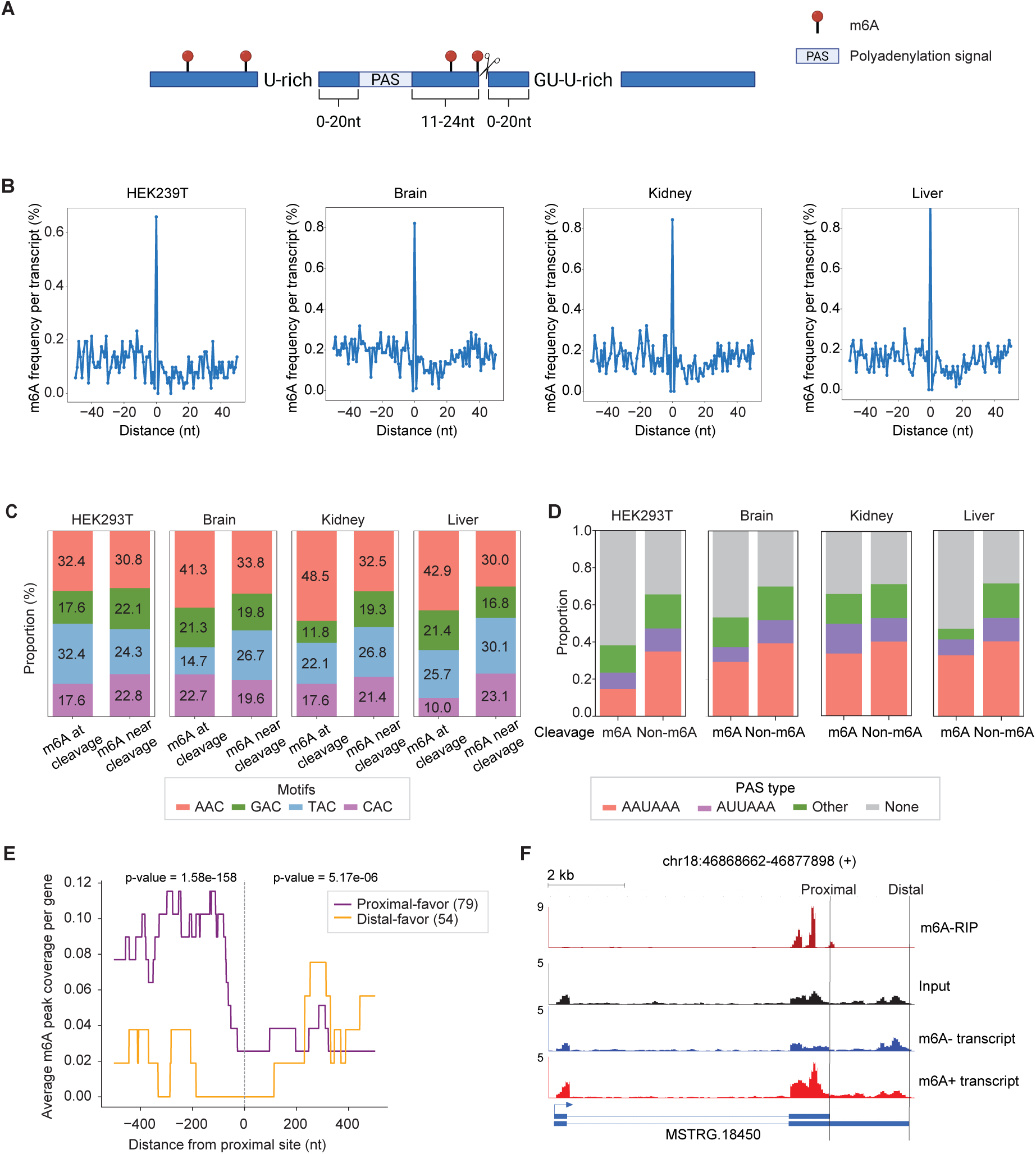
m6A is associated with non-canonical cleavage sites and alternative polyadenylation in lncRNAs. (A) Schematic diagram of a polyA site, showing the polyadenylation signal (PAS), cleavage position, and flanking sequence elements. (B) m6A site frequency within 50 nucleotides upstream and downstream of cleavage sites in lncRNAs from HEK293T cells, brain, kidney, and liver. Position 0 denotes the cleavage site. (C) Proportions of four types of m6A NAC motifs at the cleavage position and within the surrounding +/-50nt region. (D) Proportions of canonical and non-canonical polyadenylation signal (PAS) types at m6A and non-m6A cleavage sites in HEK293T cells, brain, kidney, and liver. Canonical PAS are defined as AAUAAA and AUUAAA; other motifs and sites lacking an annotated PAS are classified as non-canonical. P-values from Fisher’s exact test are 0.0145, 0.0273, 0.467, and 0.055 respectively. (E) m6A-peak coverage surrounding +/-500nt of the proximal poly(A) site in H1-ESC cells, grouped by lncRNAs with proximal-site-favored or distal-site-favored APA usage. Upstream and downstream comparisons were assessed separately using two-sided Mann–Whitney U tests. (F) Genome-browser tracks showing examples of proximal APA usage in m6A-methylated lncRNA transcripts.

In lncRNAs, we found that m6A sites are enriched at the cleavage site of lncRNAs in HEK293T cells and in human multiple tissues, rising several-fold above the flanking background **(Figure 4B**), similarly with the pattern observed in mRNA **(Figure S4A)**. Compared with m6A sites in the surrounding ±50-nt window, the motif of m6A sites at the cleavage position were more likely in an AAC context in all four samples (32.4%–48.5% versus 30.0%–33.8%; **Figure 4C**).

We next asked whether m6A-marked cleavage sites differ in their PAS. In lncRNAs, cleavage sites carrying m6A were less likely than unmethylated sites to have a canonical PAS (AAUAAA or AUUAAA) within approximately 40 nt upstream and more often lacked any recognizable PAS hexamer (**Figure 4D**). This difference was significant in HEK293T cells (P = 0.0145) and brain (P = 0.0273), showed the same trend in liver (P = 0.055), and was not significant in kidney (P = 0.467; two-sided Fisher’s exact test). In mRNAs, the same difference was larger and significant in all four samples (P = 4.77 × 10^−23^ to 3.95 × 10^−2^; **Figure S4B**). Thus, m6A sites at lncRNA 3′ ends are preferentially associated with cleavage sites that lack a canonical PAS, extending to lncRNAs an association previously described for mRNAs.^38^

Finally, we asked whether m6A-modified lncRNAs preferentially use proximal or distal polyA sites within individual lncRNA genes. Because m6A-LAIC-seq separates methylated from unmethylated transcripts of the same gene, it allows isoform usage to be compared between the two populations.^30^ For lncRNAs with tandem poly(A) sites in the same terminal exon, we compared the relative usage of proximal and distal sites between the m6A-positive and m6A-negative lncRNAs of H1-ESCs **(STAR Methods)**. We found 79 m6A-postive lncRNA transcripts with m6A near their 3’ end terminal preferentially used the proximal site (proximal-favoring; FDR < 0.05), and in 54 m6A-negative lncRNA transcripts, they preferentially used the distal site (distal-favoring; FDR < 0.05). Gene-level analysis of m6A-seq profiles around proximal poly(A) sites further showed significantly higher m6A enrichment upstream of these sites in proximal-favoring lncRNAs than in distal-favoring lncRNAs **(Figure 4E).** In contrast, m6A enrichment downstream of the proximal poly(A) site did not differ noticeably between the two groups **(Figure 4E)**. For example, the lncRNA MSTRG.18450 carries an m6A peak immediately upstream of its proximal poly(A) site; its m6A-positive transcripts predominantly terminate at the proximal site, whereas its m6A-negative transcripts extend to the distal site (**Figure 4F**).

## Discussion

Our integrative analysis links the distinct m6A landscape of lncRNAs to sequence features and RNA processing architecture. TE-derived sequences, exon organization, and 3′-end contexts are associated with methylation patterns. The response to Y14 depletion further supports a role for the EJC-associated constraint on m6A deposition in lncRNA internal exons. Together, these findings suggest that the broader distribution of m6A in lncRNAs may reflect shared deposition rules acting on differences in motif availability and transcript architecture, without necessarily requiring a lncRNA-specific targeting mechanism.

m6A-LAIC-seq analysis of genes expressed in GM12878 and H1-ESC cells revealed that lncRNAs tend to have higher gene-level methylated RNA fractions than mRNAs. This indicates that a larger proportion of RNA molecules from these lncRNA genes are methylated but does not necessarily imply higher methylation occupancy at individual sites or a greater absolute abundance of methylated RNA. Indeed, the generally higher expression levels of mRNAs may result in a higher absolute abundance of methylated molecules despite their lower methylated fractions.^43^ Whereas m6A-LAIC-seq measures gene-level methylated RNA fractions, m6A-REF-seq maps detectable m6A sites at single-base resolution. Together, these approaches provide complementary information about the prevalence of methylated transcripts and the positions m6A sites.

The TE-associated findings highlight a distinction between motif availability and detectable modification. TE-positive lncRNAs exhibit higher m6A frequencies across relative transcript positions and contain more NAC motifs than TE-negative lncRNAs, yet have a lower ratio of detected m6A sites to candidate NAC motifs. Thus, greater motif abundance does not necessarily translate into a higher proportion of motifs being detectably methylated.

Local RNA structure may contribute to the lower m6A/NAC ratio in TE-positive lncRNAs. Our observation that m6A is less frequent in stem than in non-stem regions supports a model in which base pairing limits access to potential methylation sites, consistent with previous evidence that RNA secondary structure influences m6A deposition.^44^ TE-derived sequences may contribute to such structural constraints through intramolecular base pairing. TE-positive lncRNAs also exhibited longer reverse complementary alignment within the transcript sequence and more negative predicted minimum free energies (MFE), although MFE difference may partly reflect their greater transcript lengths. Together, these findings support a model in which TE-derived sequences provide potential m6A motifs, while local RNA structure may limit the methylation of a subset of these sites.

The negative correlation between the m6A/NAC ratio and exon number links lncRNA methylation to exon architecture. Previous studies have shown that the exon junction complex (EJC) restricts m6A deposition near exon junctions in mRNAs.^32,34–36^ Our comparison of the same exons across control and Y14-depleted conditions revealed a greater relative increase in the proportion of m6A peak-containing short internal exons following Y14 depletion, supporting a similar constraint in lncRNAs. However, most short internal exons still lacked detectable m6A peak after Y14 depletion, suggesting that additional factors may limit their methylation.

Our analysis revealed m6A enrichment near lncRNA cleavage sites and a higher frequency of non-canonical polyadenylation signals among lncRNAs with m6A-marked cleavage site. This association raises the possibility that m6A plays an auxiliary role in lncRNA 3’-end processing, particularly in transcripts with non-canonical PAS. Such a role could involve enhanced access of processing factors to local RNA elements through m6A-induced structural changes, or interactions between m6A reader proteins and the 3′-end processing machinery.^12,45^ Conversely, the recruitment of processing factors or transcriptional dynamics may influence m6A deposition near cleavage sites.

Our APA analysis revealed an association between m6A methylation and alternative polyadenylation site usage in lncRNAs. In proximal-favoring genes, proximal isoforms were preferentially represented in the m6A-positive fraction, and an independent m6A-seq analysis revealed enrichment upstream of proximal poly(A) sites. In contrast, distal-favoring genes exhibited m6A enrichment within the extended regions. Together, these findings suggest that the positional distribution of m6A is associated with proximal and distal poly(A) site usage. However, whether m6A directly influences cleavage-site selection or instead reflects differences in methylation or stability among alternative isoforms remains unclear.

Our findings can be considered in the context of the recently proposed m6A RNA-surveillance model, in which splicing-associated protection limits methylation near exon junctions, whereas RNAs lacking such protection may accumulate m6A modifications and undergo reader-dependent decay.^46^ Our observation suggests that similar architectural constraints may operate in lncRNAs, potentially linking RNA processing history to m6A deposition and subsequent RNA turnover. However, whether these modifications promote the degradation of individual lncRNAs remains to be determined. Beyond RNA surveillance, m6A modification can also contribute to lncRNA regulatory functions, including XIST-mediated transcriptional silencing and MALAT1-mediated regulation of nuclear speckle organization.^22,47^ These examples suggest that the functional consequences of lncRNA methylation may depend on m6A site location, RNA localization, and the availability of specific m6A reader proteins.

Our site maps and integrative analyses identify RNA sequence and architectures features associated with m6A methylation in lncRNAs and provide a framework for investigating their functional consequences of lncRNA methylation. Future studies integrating site-specific perturbations with measurements of RNA processing and stability will help clarify the casual relationship among transcript architecture, m6A deposition and lncRNA fate.

## Limitations of the study

Our study integrates publicly available datasets generated using different assays and biological systems. The m6A-LAIC-seq and m6A-REF-seq results therefore provide complementary information about lncRNA methylation rather than directly matched measurements from the same biological samples. m6A-REF-seq relies on MazF-mediated cleavage at ACA motifs and therefore does not detect m6A sites in other sequence contexts, including ACC and ACU. Detection also depends on sequencing coverage and MazF cleavage efficiency, which can be influenced by local RNA secondary structure. Additional limitations include the selection of the longest isoform to represent each gene, different expression or count thresholds for lncRNAs and mRNAs. Poly(A)-enriched sampling limits the representation of non-polyadenylated lncRNAs, restricting the generalizability of our findings to the broader lncRNA transcriptome. Most analyses are observational. Although the published Y14-depletion experiment provides perturbation-based evidence, it does not fully distinguish direct effects on m6A deposition from indirect consequences of Y14 depletion. We did not directly measure the effects of m6A on lncRNA function, decay, or processing kinetics. Establishing these causal relationships will require targeted functional studies.

## STAR Methods

| REAGENT OR RESOURCE | SOURCE | IDENTIFIER |
|---|---|---|
| **Deposited data** |  |  |
| m6A-REF-seq: HEK293T and human brain, kidney, liver | Zhang et al.<sup>31</sup> | GEO: GSE125240 |
| m6A-LAIC-seq: H1-ESC and GM12878 | Molinie et al.<sup>30</sup> | GEO: GSE66086 |
| Y14-depletion m6A-IP data | Uzonyi et al.; Luo et al.<sup>32,35</sup> | GEO: GSE204980 |
| H1-ESC m6A-seq | Batista et al.<sup>13</sup> | GEO: GSE52600 |
| **Annotation and processed analysis outputs** |  |  |
| Human reference genome GRCh38/hg38 | Genome Reference Consortium | https://www.ncbi.nlm.nih.gov/datasets/genome/GCF_000001405.26/ |
| LncBook 2.0 | Li et al.<sup>48</sup> | doi:10.1093/nar/gkac999 |
| Comprehensive gene annotation (v29) | GENCODE | https://www.gencodegenes.org/human/release_29.html |
| RepeatMasker TE annotation | UCSC Genome Browser | http://genome.ucsc.edu/cgi-bin/hgTables |
| PolyA_DB 3.2 | Wang et al.<sup>42</sup> | doi:10.1093/nar/gkx1000 |
| lncRNA annotation and methylation summaries | This study | Tables S1–S5 |
| Custom poly(A) site annotation generated in this study | This study | Table S6 |
| **Software and algorithms** |  |  |
| Flnc v2.0 | Li et al.<sup>49</sup> | doi:10.3390/ncrna8050070; release |
| HISAT2 2.2.1 | Kim et al.<sup>50</sup> | doi:10.1038/s41587-019-0201-4 |
| StringTie 2.2.1 | Pertea et al.<sup>51</sup> | doi:10.1038/nbt.3122 |
| FastQC 0.11.9 | Andrews | https://www.bioinformatics.babraham.ac.uk/projects/fastqc/ |
| Cutadapt 4.6 | Martin<sup>52</sup> | doi:10.14806/ej.17.1.200 |
| RNAfold/ViennaRNA 2.6.3 | Lorenz et al.<sup>53</sup> | doi:10.1186/1748-7188-6-26 |
| BEDTools 2.26.0 | Quinlan and Hal<sup>54</sup> | doi:10.1093/bioinformatics/btq033 |
| HTSeq1.99.2 | Anders et al.<sup>55</sup> | doi:10.1093/bioinformatics/btu638 |
| BLAST+ 2.17.0 | Camacho et al.<sup>56</sup> | doi:10.1186/1471-2105-10−421 |
| UCSC LiftOver | UCSC | https://genome.ucsc.edu/cgi-bin/hgLiftOver |
| APAtrap | Ye et al.<sup>57</sup> | doi:10.1093/bioinformatics/bty029 |
| Custom rMATS-based APA analysis | Shen et al.<sup>58</sup>; this study | doi:10.1073/pnas.1419161111; implementation |
| exomePeak2 3.18 | Wei<sup>59</sup> | Bioconductor 3.18 |
| Statistical/figure generation environment | This study | R/Python packages and environment |

### Data and code availability

Public sequencing data analyzed in this study are available from the Gene Expression Omnibus (GEO) under accessions GSE125240 (m6A-REF-seq), GSE66086 (m6A-LAIC-seq), GSE204980 (EJC/Y14-depletion m6A-IP), and GSE52600 (H1-ESC m6A-seq). The sample identifiers, fractions or treatment conditions, and replicate assignments used for each analysis are listed in STAR Methods. The lncRNA annotation, gene based m6A-level estimates, site calls, TE classifications, internal-exon summaries and custom poly(A) site annotation are provided in Tables S1–S6. Additional information required to reanalyze the reported data is available from the lead contact upon request.

### Construction of the lncRNA reference used in this study

To establish a comprehensive lncRNA reference for downstream analyses, we integrated annotated lncRNAs with sample-specific novel lncRNAs. Annotated lncRNA transcripts were obtained from LncBook 2.0^48^ in GTF format and converted to BED format. To assess the reliability of the annotated lncRNAs, Flnc v2.0^49^ was applied in single-input mode using the BED-formatted annotations (Flnc.py single-f bed-u Lncbookv2.0_annotation.bed) to filter out potential transcript fragments and transcriptional noise.

To identify novel lncRNAs in individual samples, adapter-trimmed paired-end RNA-seq reads from the m6A-REF-seq dataset (GEO: GSE125240) were processed using Flnc with the default random forest classification model (-m rf) and the second-strand library orientation (-s second). Finally, the filtered annotated lncRNAs and newly identified sample-specific lncRNAs were combined, and redundant transcripts were removed to generate the final lncRNA reference annotation (Table S1), which was used for subsequent analyses.

### m6A level estimates from m6A-LAIC-seq

lncRNA gene expressed in H1-ESC and GM12878 sample were identified using Flnc v2.0^49^ from input, m6A-positive (eluate) and m6A-negative (supernatant) of m6A-LAIC-seq data (GEO Accession number: GSE66086) with ‘-m rf-s first ‘options. Identified novel lncRNA gene were integrated to reference gene annotation (Gencode v29), followed by quantification of gene abundance using HTSeq.^55^

To calculate m6A levels for each lncRNA/mRNA gene, we used the linear regression of ERCC RNA values provided in the original m6A-LAIC-seq paper (GM12878_sp1 = 1.585, GM12878_sp2= 0.944, H1_sp1 = 1.412, H1_sp2= 1.468).^30^ To obtain reliable m6A level for each mRNA and lncRNA gene, we required eluate counts ≥50 or supernatant counts ≥50 in both replicates in mRNA and eluate counts ≥5 or supernatant counts ≥5 in both replicates in lncRNA, as the median lncRNA expression level is approximately tenfold lower than the median protein-coding mRNA level.^43^ The m6A levels of each gene were calculated using eluate counts (E), supernatant counts (S), and the residual of ERCC regression (R) according to the formula: m6A level = E / (E + S × 2^R).

### m6A-REF-seq read processing, alignment, and expression quantification

Raw paired-end m6A-REF-seq reads (GEO: GSE125240) were assessed for sequencing quality using FastQC,^60^ and adapters sequences were trimmed using cutadapt,^52^ retaining reads with a minimum length of 15 nt. Trimmed reads were aligned to the human reference genome (GRCh38/hg38) using HISAT2^50^ with default parameters. Transcript abundance was quantified using StringTie^51^ in expression-estimation mode (-e), guided by the combined transcript annotation (-G). Gene-level abundance estimates were generated using the-A option. Genes with FPKM ≥ 0.1for lncRNAs and FPKM ≥1 for mRNAs were considered expressed. For transcript-level analyses, the longest annotated transcript of each expressed gene was selected as its representative isoform.

### Single-base m6A detection from m6A-REF-seq

Single-nucleotide resolution m6A sites were identified by adapting a previously published m6A-REF-seq computational pipeline.^31^ This method exploits the sensitivity of MazF-mediated cleavage at ACA motifs to methylation of the first adenosine. Reads aligned to ACA motifs on the positive strand or the corresponding reverse-complement TGT motifs on the negative strand were classified according to whether they spanned the motif internally or terminated at the expected cleavage position. An internal motif was required to be located more than 15 nt from both ends of the aligned read, and candidate sites were required to have at least 10 supporting reads containing an internal motif. Candidate sites containing repeated or consecutive ACA motifs were excluded.

For each candidate site, the relative fraction of reads spanning the uncleaved motif was calculated as r = I/(I + C), where I represents the number of reads containing an internal ACA motif and C represents the number of reads supporting cleavage at the corresponding motif. RNA secondary structure was evaluated using RNAfold from the ViennaRNA package^53^ to identify candidate sites potentially affected by intramolecular base pairing. Sites with high predicted pairing probabilities were excluded. To further validate m6A candidate sites, FTO-demethylated RNA samples were used as negative controls. Candidate sites exhibiting a decrease of at least 0.10 in the uncleaved-read fraction following FTO treatment were retained as putative m6A sites. Site calling was performed independently for each HEK293T biological replicate, and sites detected in at least two replicates were retained for downstream analyses (Table S3).

### Transcript-level distribution and frequency analysis of m6A sites

To characterize the transcript-level distribution of m6A sites, the longest transcript of each expressed gene was selected as the representative isoform. For metagene analysis, lncRNA transcripts were divided into 100 equal-length bins. Each mRNA transcripts were partitioned into the 5′ UTR, CDS, and 3′ UTR, which were divided into 25, 50, and 25 equal-length bins, respectively. For lncRNAs, m6A frequency was calculated as the number of m6A sites per bin divided by the number of expressed lncRNA transcripts. For mRNAs, m6A frequency was calculated as the number of m6A sites per bin divided by the number of expressed mRNA transcripts containing the corresponding region. Length-normalized m6A frequency per kilobase were calculated using the same binning schemes, with site counts in each bin normalized by the corresponding bin length and the number of expressed lncRNA transcripts or mRNA transcripts containing the corresponding region and multiplied by 1,000.

To visualize the distribution of m6A sites across individual transcripts, heatmaps were generated for transcripts containing at least one identified m6A site. LncRNA transcripts were divided into 20 equal-length bins, while the 5′ UTR, CDS, and 3′ UTR of mRNA transcripts were divided into 5, 10, and 5 bins, respectively. The number of m6A sites assigned to each bin was used to construct the heatmaps. Transcript-level m6A frequency was calculated as the number of identified m6A sites divided by the spliced transcript length.

### Exon-based positional analysis of m6A distribution

The longest representative transcripts of expressed lncRNA and mRNA genes were stratified into single-exon, two-exon, and ≥3-exon groups. The same exon-based binning and normalization schemes were applied to both RNA classes in HEK293T cells and human brain, kidney, and liver tissues. Single-exon transcripts were divided into three equal-length regions, designated first, internal, and last, each comprising 30 bins. For two-exon transcripts, the first and last exons were each divided into 45 equal-length bins. For transcripts with three or more exons, the first, internal, and last exon were each represented by 30 equal-length bins. Internal exons were defined as all exons between the first and last exon in the 5’ to 3’ direction of transcription. m6A site counts were normalized by the corresponding bin length and the number of expressed transcripts contributing to each region and multiplied by 1,000 to obtain length-normalized m6A frequency per kilobase. Mean regional m6A frequency was calculated by averaging the normalized bin values within each region or exon. For single-exon transcripts, fold changes were calculated for the internal versus first, last versus internal, and last versus first regions. For two-exon transcripts, the last exon was compared with the first exon. For transcripts with three or more exons, fold changes were calculated by dividing the mean m6A density of each exon (first, internal and last) by that of the first exon, yielding a reference value of 1 for the first exon.

### Transposable element (TE) annotation and classification of TE+ and TE− transcripts

To characterize the distribution of transposable elements (TEs) across lncRNA and mRNA transcripts, TE annotations were obtained from the UCSC RepeatMasker track. The annotations were filtered to retain major TE classes, including LINE, SINE, DNA, LTR, and Retroposon elements, while non-TE annotations, such as low-complexity regions, satellites, simple repeats, and ncRNAs, were excluded. The retained annotations were converted to BED format and intersected with the selected transcript models using BEDTools intersect. Transcripts containing at least one retained TE annotation overlapping an exonic region were classified as TE-positive (TE+), whereas transcripts without any exonic TE overlap were classified as TE-negative (TE−) (Table S4). TE positions were mapped to normalized transcript coordinates using the same binning strategy applied to the m6A metagene profiles. The distributions of m6A frequency and NAC motifs were analyzed separately for TE+ and TE− transcripts using the same binning and length-normalization approach described above.

### RNA secondary structure prediction and classification of m6A sties

To investigate the potential influence of TEs on RNA secondary structure and m6A distribution, full-length lncRNA transcript sequences and mRNA sequences comprising the 3′ UTR and the 100-nt upstream region were extracted from TE+ and TE− transcripts. Minimum free energy (MFE) values and base-pairing probabilities were calculated for each sequence using RNAfold from the ViennaRNA package.^53^

To characterize the local structural context of m6A sites in both TE+ and TE− transcripts, nucleotide positions were classified as stem or non-stem regions based on the predicted secondary structures represented in dot-bracket notation. A position was classified as a stem region if the nucleotide itself and both immediately adjacent nucleotides (−1 and +1 nt) were base-paired. All remaining positions were classified as non-stem regions. The number of m6A sites were adjusted by the total sequence length of each structural region to plot the proportion of m6A site in stem versus non-stem regions.

### Analysis of intramolecular sequence complementarity using BLASTN

To assess the potential for intramolecular sequence complementarity in TE+ and TE− transcripts, full-length lncRNA sequences and mRNA sequences comprising the 3′ UTR and the 100-nt region upstream of the stop codon were extracted. Sequence alignments were performed using NCBI BLASTN (v2.17.0+)^56^ with relaxed parameters (-word_size 7,-gapopen 3,-gapextend 1,-reward 1,-penalty-1, and-evalue 100) to identify potential complementary regions. Intramolecular reverse-complement alignments were identified by retaining hits with matching query and subject sequence identifiers (qseqid = sseqid) and a minus-strand orientation (sstrand = minus).

### Correlation between m6A/NAC motif ratio and exon number

To investigate the relationship between m6A frequency and exon number, the ratio of identified m6A sites to NAC motifs (m6A/NAC ratio) was calculated for the longest transcript of each expressed lncRNA and mRNA gene. The m6A/NAC ratio was plotted against the number of exons per transcript, and linear regression analysis was performed to evaluate their association. The regression slope and corresponding p-value were calculated.

### Analysis of m6A enrichment in lncRNA internal exons following EJC depletion

To evaluate the influence of the exon junction complex (EJC) on m6A deposition, we used previously processed m6A peak data provided by Luo et al.,^32^ derived from m6A-IP and input sequencing data from control and Y14-depleted HEK293T cells (two biological replicates per condition; GEO: GSE204980).^35^ Y14, a core component of the EJC, was depleted using the dTAG degron system. Peaks with a fold enrichment greater than 1 (m6A-IP relative to input) were retained. LncRNA transcript models were obtained from the Flnc-based annotation, with the longest expressed transcript selected for each gene in HEK293T cells. Transcripts containing at least three exons were retained, and the first and last exons were excluded. The remaining exons were defined as internal exons and classified as short (≤200 nt) or long (>200 nt) based on their length. For each condition, we quantified the number of internal exons containing at least one NAC motif and the number overlapping at least one m6A peak (Table S5). Positional m6A enrichment was calculated as the fraction of internal exons overlapping an m6A peak within each 10-nt bin, extending up to 180 nt from the 5′ end of the exon. Only exons extending to the corresponding bin were included in the calculation. This analysis was performed separately for each biological replicate, and the mean of the two replicates was plotted.

### Construction of a comprehensive poly(A) site reference

Reference human cleavage and polyadenylation sites were obtained from PolyA_DB (v3.2)^42^, and duplicate sites were removed. Poly(A) sites were classified as mRNA-or lncRNA-associated based on their genomic annotations and RefSeq accession prefixes. Genomic coordinates were converted from hg19 to hg38 using the UCSC LiftOver tool. To identify additional poly(A) sites from RNA-seq data, APAtrap^57^ was applied using the ‘identifyDistal3UTR’ and ‘predictAP’ modules. The ‘identifyDistal3UTR’ module was used to refine annotated 3′ UTRs and identify potential 3′ UTR extensions. For HEK293T and human tissue datasets, the module was run using a scanning window of 20 nt (-w 20), a maximum extension length of 15,000 nt (-e 15000), a minimum relative distal-end coverage threshold of 0.01 (-c 0.01), and a minimum proportion of covered nucleotides per scanning window of 0.5 (-p 0.5). For H1 ESC data used in the alternative polyadenylation analysis, the corresponding parameters were set to-w 50,-c 0.05, and-p 0.2, while unspecified parameters were left at their default values. Subsequently, the ‘predictAP’ module was used to identify candidate poly(A) sites with a minimum average coverage of 10 (-c 10), a minimum degree of coverage variation between adjacent APA sites of 0.1 (-d 0.1), a minimum distance of 20 nt between predicted APA sites (-a 20), and a scanning window of 20 nt (-w 20). Finally, predicted poly(A) sites that did not overlap with PolyA_DB reference sites were combined with the reference annotations to generate a comprehensive poly(A) site set (Table S6).

### Association of m6A with lncRNA cleavage and polyadenylation

To characterize the distribution of m6A sites around cleavage and polyadenylation sites, genomic regions spanning 50 nt upstream and downstream of each cleavage site were generated using BEDTools slop (-l 50-r 50-s). Identified m6A sites were intersected with these regions, and positional m6A frequency was calculated as the proportion of transcripts containing an m6A site at each position relative to the cleavage site. To examine m6A motif composition, m6A sites were classified according to their positions at the cleavage site or within the surrounding ±50-nt region, and the proportions of individual m6A motifs were calculated for each region. To investigate the association between m6A modification and polyadenylation signal (PAS) usage, poly(A) sites were classified as m6A-associated or non-m6A-associated based on the presence or absence of an overlapping m6A site. The 40-nt sequences upstream of each poly(A) site were extracted and screened for canonical (AAUAAA and AUUAAA) and non-canonical (AGUAAA, UAUAAA, CAUAAA, GAUAAA, AAAAAA, AAUAUA, AAUACA, AAUAGA, AAAAAG, and ACUAAA) PAS hexamers. Sites without a match to these hexamers were classified as “None”. Differences in the proportions of canonical and non-canonical PAS categories between m6A-associated and non-m6A-associated poly(A) sites were assessed using a two-sided Fisher’s exact test.

### Differential tandem polyadenylation analysis and m6A peak coverage profiling

To identify novel poly(A) sites from H1 ESC RNA-seq data, the ‘identifyDistal3UTR’ and ‘predictAP’ modules of APAtrap^57^ were applied to two biological replicates of the input fraction from the m6A-LAIC-seq dataset. The predicted poly(A) sites were combined with reference sites from PolyA_DB (v3.2)^42^, with redundant sites removed, to generate a custom poly(A) site annotation. To investigate differential poly(A) site usage between m6A-positive and m6A-negative RNA fractions, poly(A) sites from this custom annotation were mapped to the terminal exons of expressed lncRNA genes. For transcripts with predicted 3′ UTR extensions identified by ‘identifyDistal3UTR’, the terminal exon boundaries were extended accordingly. Tandem APA events were defined by proximal and distal poly(A) sites located within the same terminal exon. For transcripts containing multiple poly(A) sites within a terminal exon, proximal–distal site pairs were constructed to define candidate tandem APA events. The region shared by both isoforms was designated as the common terminal-exon segment, whereas the region between the proximal and distal poly(A) sites was designated as the extended terminal-exon segment. Only tandem APA pairs with both common and extended segments longer than 100 nt were retained. For each tandem APA pair, RNA-seq reads mapping to the common and extended terminal-exon segments were quantified separately for each biological replicate of the m6A-positive and m6A-negative fractions. Read counts were normalized by the corresponding segment lengths and used to estimate relative proximal and distal poly(A) site usage. The statistical framework implemented in rMATS^58^ was adapted to assess differences in APA usage between the two fractions. Tandem APA pairs with fewer than two reads in total across the common and extended segments in any replicate were excluded.

Tandem APA pairs with a false discovery rate (FDR) < 0.05 were considered to exhibit differential APA usage. For genes containing multiple significant tandem APA pairs, the pair with the lowest FDR was selected for downstream analyses. Genes were classified into three groups based on the direction and significance of differential APA usage: Group 1, preferential proximal poly(A) site usage in the m6A-positive fraction; Group 2, preferential distal poly(A) site usage in the m6A-positive fraction; and Group 3, no significant difference in APA usage (FDR ≥ 0.05).

To investigate the relationship between m6A enrichment and differential APA usage, m6A peaks were identified from the H1 ESC m6A-IP eluate and input samples (GEO: GSE52600)^13^ using exomePeak2.^59^ m6A peak coverage was profiled around proximal poly(A) sites in Group 1 and Group 2 genes. For each gene, the transcript isoform containing the extended terminal-exon segment was used to calculate m6A peak coverage within a 500-nt window centered on the proximal cleavage site. Transcripts lacking sufficient flanking sequence to span the entire window were excluded. To account for differences in terminal-exon architecture, the longest annotated terminal exon among transcript isoforms was selected for each gene.

## Resource availability

### Lead contact

Requests for further information and resources should be directed to and will be fulfilled by the lead contact, Chan Zhou.

### Materials availability

This study did not generate new unique reagents.

### Data and code availability

This paper analyzes existing, publicly available data. Accession numbers are listed in the key resources table (GEO: GSE125240, GSE66086, GSE204980, and GSE52600). The lncRNA annotation, single-base m6A sites, and other processed data generated in this study are provided in Tables S1–S6. Any additional information required to reanalyze the data reported in this paper is available from the first contact upon request as of the date of publication.

## Acknowledgments

We thank Dr. Shengdong Ke lab and his laboratory for providing the m6A-IP data of Y14 control and Y14-depletion conditions, which result were shown in their previous publication.^32^ We also thank Dr. Zhiping Weng, Dr. Alan C. Mullen, Dr. Athma Pai, and Dr. Tianxiong Yu of UMass Chan Medical School for insightful suggestions on this study. This work was supported by start-up funds from UMass Chan Medical School.

## Author contributions

EK and CZ conceived the study. EK and CZ designed the study. EK performed most of data analyses and ZL performed the initial m6A-LAIC-seq data analysis for lncRNAs. EK and CZ interpreted the results. EK and CZ drafted the manuscript, with input and revisions from ZL.

## Declaration of generative AI and AI-assisted technologies in the writing process

During the preparation of this work, the authors used ChatGPT to correct grammar and spelling, and to improve the English language readability. After using this tool, the authors reviewed and edited the output as needed and take full responsibility for the content of the published article.

