## Supplemental Figures for "Single-base mapping of m6A in lncRNAs reveals a distinct landscape linked to transposable elements and RNA processing"

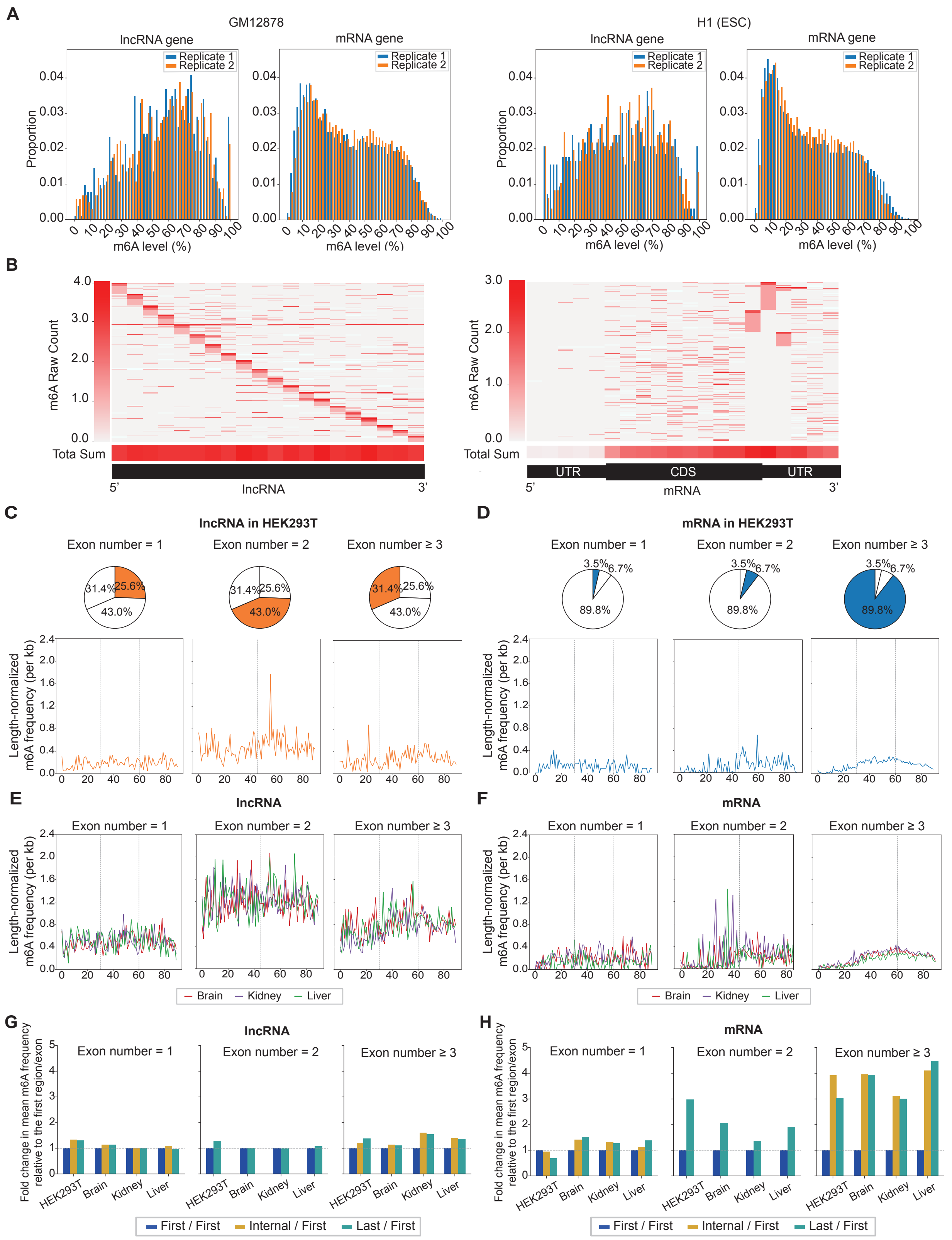


**Figure S1. Single-base-resolution mapping reveals distinct m6A landscapes in lncRNAs and mRNAs (related to Figure 1). (A)** Distributions of transcript-level m6A levels for lncRNAs and mRNAs in GM12878 and H1 ESCs. Replicates 1 and 2 are shown in blue and orange, respectively. **(B)** Heatmap of m6A sites across individual lncRNA and mRNA transcripts. Each rows represents an individual transcript. lncRNA transcripts were ranked from the highest to the lowest m6A enrichment within each bin. mRNA transcripts were ranked according to the m6A enrichment at the last bin of CDS and the first bin of 3'UTR. The bottom 'Total Sum' tracks show the summed m6A signal across all transcripts at each bin. **(C)** Length-normalized m6A frequency per kilobase across single-exon, two-exon, and ≥3-exon lncRNA transcripts in HEK293T cells. Single-exon transcripts were divided into three equal regions, referred to as the first, internal, and last region. For two-exon transcripts, the first and last exons were each represented by 45 equal-length bins. For transcripts containing three or more exons, the first, internal, and last exon regions were each represented by 30 equal-length bins. m6A site counts were normalized by the corresponding bin length and the number of expressed transcripts contributing to each region and multiplied by 1,000. **(D)** Length-normalized m6A frequency per kilobase across single-exon, two-exon, and ≥3-exon mRNA transcripts in HEK293T cells, using the same binning and normalization scheme as in (C). **(E)** Length-normalized m6A frequency per kilobase across single-exon, two-exon, and ≥3-exon lncRNA transcripts in multiple human tissues, using the same binning and normalization scheme as in (C). **(F)** Length-normalized m6A frequency per kilobase across single-exon, two-exon, and ≥3-exon mRNA transcripts in multiple human tissues, using the same binning and normalization scheme as in (C). **(G)** Fold changes in mean length-normalized m6A frequency across lncRNA transcript regions relative to the corresponding first exon or first transcript region. For two-exon transcripts, fold changes were calculated for the last versus first exons. For ≥3-exon transcripts, fold changes were calculated for internal versus first, last versus internal, and last versus first exon regions. **(H)** Fold changes in mean length-normalized m6A frequency across mRNA transcript regions, calculated using the same comparisons as in (G).

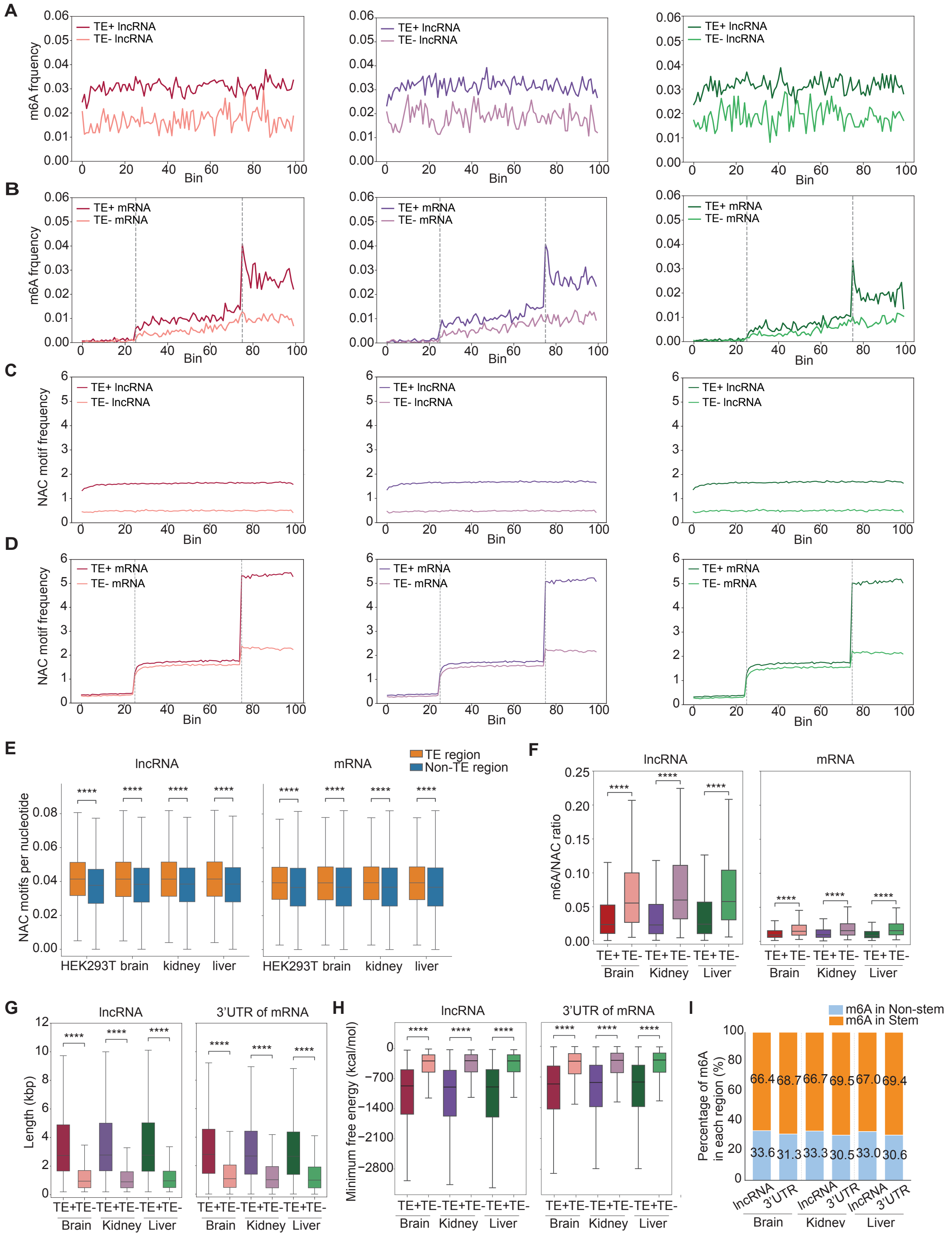


**Figure S2. Transposable element-derived sequences and local RNA structure may contribute to distinct m6A patterns between lncRNAs and mRNAs across multiple human tissues (related Figure 2). (A)** Distribution of m6A frequency across TE-positive (TE+) and TE-negative (TE−) lncRNAs in brain (left), kidney (middle), and liver (right). **(B)** Distribution of m6A frequency across TE-positive (TE+) and TE-negative (TE−) mRNAs in brain (left), kidney (middle), and liver (right). **(C)** Distribution of NAC motif frequency across TE-positive and TE-negative lncRNAs in brain (left), kidney (middle), and liver (right). N denotes any nucleotide. **(D)** Distribution of NAC motif frequency across TE-positive and TE-negative mRNAs in brain (left), kidney (middle), and liver (right). **(E)** Number of NAC motifs normalized by the total sequence length of TE-overlapping (TE-region) and non-TE regions in lncRNA and mRNA. **(F)** m6A/RAC ratios in TE-positive and TE-negative lncRNAs and mRNAs. **(G)** Length distributions of TE-positive and TE-negative lncRNAs and mRNA 3′ UTRs**. (H)** Minimum free energy (MFE) of TE-positive and TE-negative lncRNAs and mRNA 3′ UTRs.m6A/RAC ratios in TE-positive and TE-negative lncRNAs and mRNAs. **(I)** Length-normalized proportion of m6A sites located in stem and non-stem regions of lncRNAs and mRNA 3′ UTRs.

For (A)-(D), darker and lighter lines represent TE-positive and TE-negative transcripts, respectively. Statistical significance was assessed using two-sided Mann–Whitney U tests for (E)–(H). ****$p \leq 1 \times 10^{-4}$.

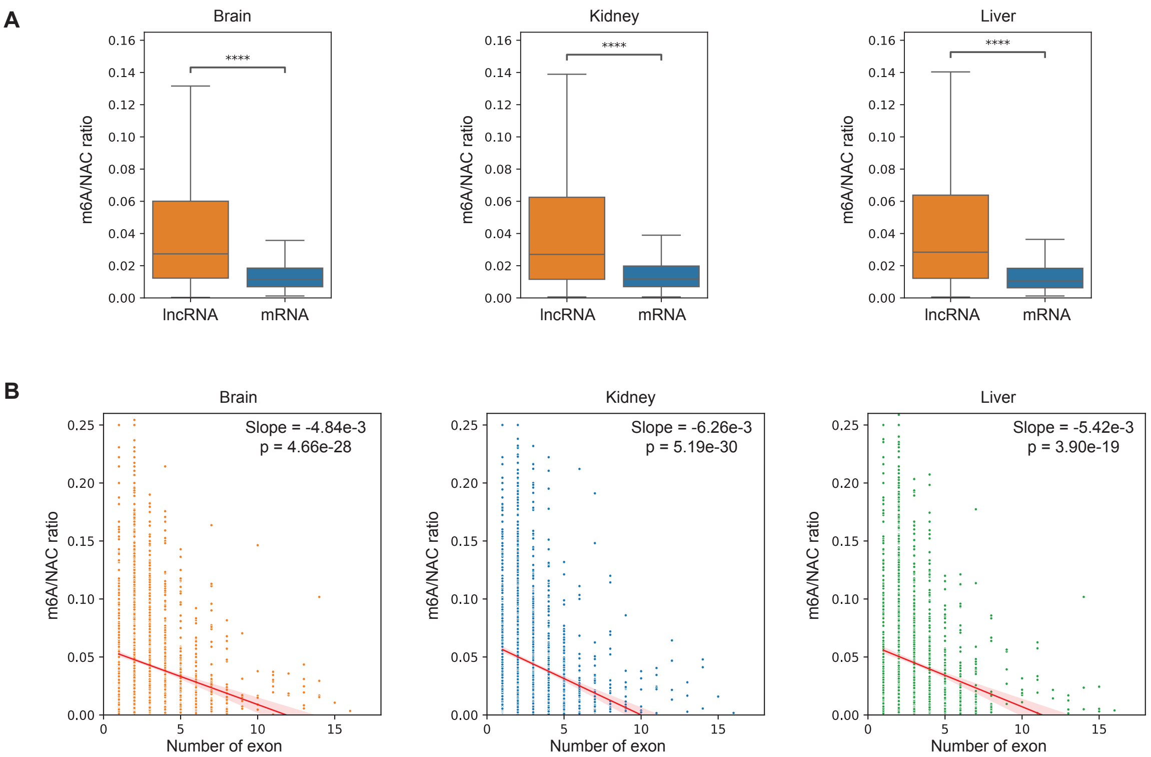


**Figure S3. m6A deposition in lncRNA is negatively correlated with exon number (related to Figure 3) (A)** m6A/RAC ratio in lncRNA and mRNAs. **** denotes p<= 1.00e-04 which were calculated with Mann-Whitney-Wilcoxon two-sided test. **(B)** The scatter plot showing the relationship m6A/RAC ratio and exon numbers for individual lncRNA transcripts. Slope and p value were calculated using the linear regression model.

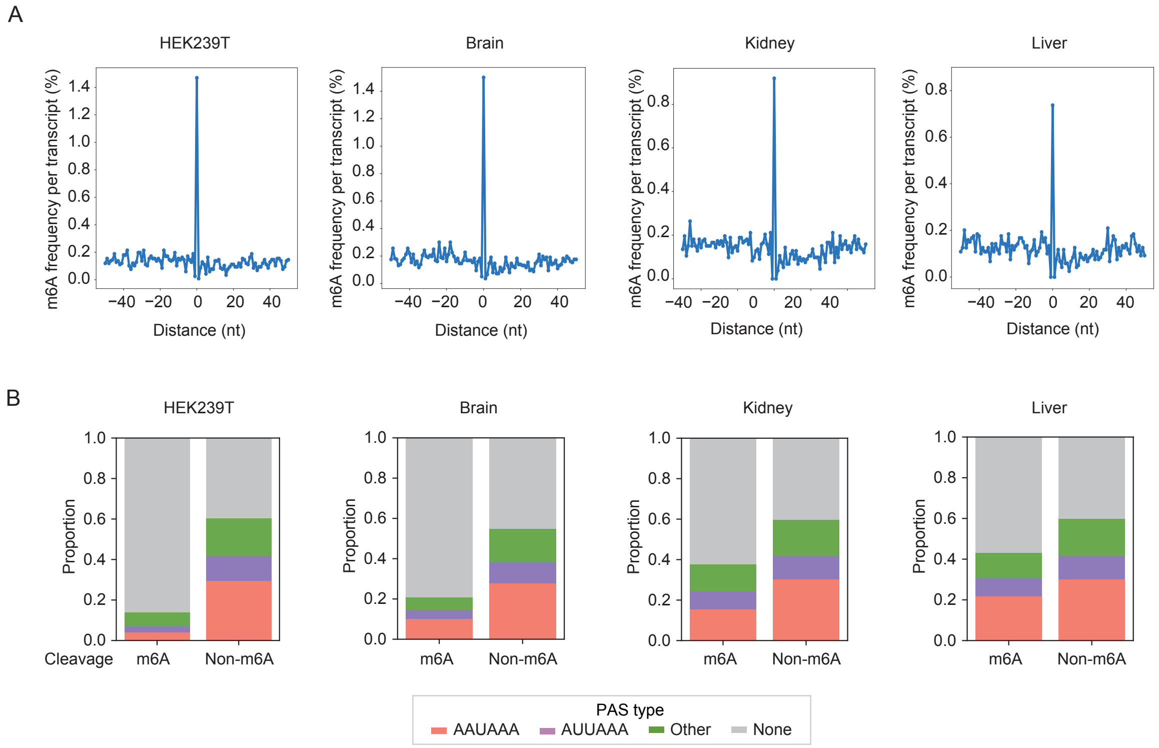


**Figure S4. m6A is associated with non-canonical cleavage sites in mRNAs (related Figure 4). (A)** m6A site frequency within 50 nucleotides upstream and downstream of cleavage sites in HEK293T cells, brain, kidney, and liver. Position 0 denotes the cleavage site. **(B)** Proportions of canonical and non-canonical polyadenylation signal (PAS) types at m6A and non-m6A cleavage sites in HEK293T cells, brain, kidney, and liver. Canonical PAS are defined as AAUAAA and AUUAAA; other motifs and sites lacking an annotated PAS are classified as non-canonical. P-values of HEK293T, brain, kidney, and liver from Fisher's exact test are 4.77e-23, 1.17e-13, 9.89e-5, and 3.95e-2 respectively.
