## Supplementary material for "Single-base mapping of m6A in lncRNAs reveals a distinct landscape linked to transposable elements and RNA processing": Information of Supplemental Tables

**Supplemental Tables S1-S6**

**Table S1: lncRNAs used in the study**

**Table S2: m6A level from m6A-LAIC-seq**

**Table S3: Single-base resolution m6A sites from m6A-REF-seq data**

**Table S4: TE region in lncRNAs and mRNAs [1]**

**Table S5: Internal exons with m6A and NAC**

**Table S6: Comprehensive polyA sites used in the study**


1. Kapusta, A., Kronenberg, Z., Lynch, V.J., Zhuo, X., Ramsay, L., Bourque, G., Yandell, M., and Feschotte, C. (2013). Transposable elements are major contributors to the origin, diversification, and regulation of vertebrate long noncoding RNAs. PLoS Genet *9*, e1003470. 10.1371/journal.pgen.1003470.
